# CRISPR/Cas9-targeted removal of unwanted sequences from small-RNA sequencing libraries

**DOI:** 10.1101/319715

**Authors:** Andrew A. Hardigan, Brian S. Roberts, Dianna E. Moore, Ryne C. Ramaker, Angela L. Jones, Richard M. Myers

**Affiliations:** HudsonAlpha Institute for Biotechnology, Huntsville, AL 35806, USA; Department of Genetics, University of Alabama at Birmingham, Birmingham, AL 35294, USA

## Abstract

In small RNA (smRNAs) sequencing studies, highly abundant molecules such as adapter dimer products and tissue-specific microRNAs (miRNAs) inhibit accurate quantification of lowly expressed species. We previously developed a method to selectively deplete highly abundant miRNAs. However, this method does not deplete adapter dimer ligation products that, unless removed by gel-separation, comprise most of the library. Here, we have adapted and modified recently described methods for CRISPR/Cas9–based Depletion of Abundant Species by Hybridization (“DASH”) to smRNA-seq, which we have termed miRNA and Adapter Dimer - DASH (MAD-DASH). In MAD-DASH, Cas9 is complexed with sgRNAs targeting adapter dimer ligation products, alongside highly expressed tissue-specific smRNAs, for cleavage *in vitro*. This process dramatically reduces (>90%) adapter dimer and targeted smRNA sequences, is multiplexable, shows minimal off-target effects, improves the quantification of lowly expressed miRNAs from human plasma and tissue derived RNA, and obviates the need for gel-separation, greatly increasing sample throughput. Additionally, the method is fully customizable to other smRNA-seq preparation methods. Like depletion of ribosomal RNA for mRNA-seq and mitochondrial DNA for ATAC-seq, our method allows for greater proportional read-depth of nontargeted sequences.

## INTRODUCTION

Small RNAs (smRNAs) are a diverse class of RNA molecules, including microRNAs (miRNAs), transfer RNAs (tRNAs), small nucleolar RNA (snoRNA), Y-RNA, and many others, that have diverse roles in biological processes (1, 2). miRNAs are particularly well-studied due to their role as post-transcriptional regulators of gene expression in many biological processes (3–5), and altered expression of smRNAs has been implicated in many disease pathologies (6–11). Consequently, there is a need for methods that can precisely and accurately measure smRNAs.

Although smRNAs are measurable with techniques such as qPCR and hybridization based methods, sequencing of smRNA libraries has distinct advantages due to its relative high-throughput and sensitive detection of numerous smRNA species (12–14). Additionally, because it allows for agnostic detection of unknown species, novel smRNAs and sequence variation of known smRNAs (such as miRNA isoMirs) can be assessed. Nevertheless, technical challenges in library preparation limit throughput and can lower library quality. In many protocols, documented ligation biases resulting from the use of ATP-turnover deficient truncated T4 RNA Ligase 2 and specific adapter sequences combined with preferential ligation of overabundant sequences can limit accuracy and make the detection of non-favored or lowly expressed smRNAs difficult (15–19). In addition, the formation of large quantities of unwanted adapter dimer ligation product necessitates time- and labor-intensive removal steps via denaturing gel-electrophoresis, as such dimers are only ~20-30 bp smaller than many desired sequences such as miRNAs.

Targeted reduction of specific sequences from sequencing libraries is frequently employed to enrich for sequences of interest, such as with rRNA- or globin-RNA reduction from mRNA-sequencing libraries. Recently, we and others have demonstrated techniques to deplete specific smRNA sequences from sequencing libraries by using blocking oligonucleotides that prevent 5’ adapter ligation, effectively preventing further incorporation of these targets in downstream library construction (20, 21). While very effective, both strategies have limitations, such as off-targets due to sequence similarity, particularly in the seed region of non-targeted miRNAs with hairpin-oligo blocking. Additionally, neither of these methods address excess adapter dimer ligation products, and thus still require the use of denaturing gel separation of library products before sequencing. Strategies using locked-nucleic acid (LNA) oligos have been employed to prevent adapter dimer formation with some limited success (22). Currently, the most effective means of adapter dimer prevention or removal during smRNA sequencing has been demonstrated with ligation free template-switching protocols or chemically modified adapters that sterically inhibit ligation of the 3’ and 5’ adapters to each other and inhibit adapter dimer reverse transcription (23, 24). Because adapter dimer formation is limited, these strategies allow for the use of SPRI-bead based size selection in place of gel separation, which greatly increases library preparation throughput. While very effective at a range of RNA input concentrations, the chemical modifications’ efficacy appears to have some adapter-sequence specificity, which, combined with reported necessity of custom reaction conditions, limits their use in other custom or commercial smRNA protocols. In addition, these methods also do not allow for the targeted removal of non-adapter dimer smRNAs, requiring additional experimental methods or greater sequencing depth to allow measurement of lowly abundant smRNAs in the background of a few highly abundant species. Considering the benefits and limitations of these approaches, a single method that can efficiently remove both unwanted smRNAs and adapter dimer from smRNA libraries in a customizable manner would be tremendously useful.

New genome- and epigenome-editing tools based on repurposing the CRISPR/Cas9 (clustered regularly interspersed short palindromic repeats / Cas9) bacterial immune system have many potential uses (25, 26). The Cas9 nuclease, when complexed with a short RNA oligonucleotide known as a single guide RNA, or sgRNA, can induce double-stranded breaks (DSBs) at specific sgRNA complementary locations. The low-cost and easily programmable nature of the CRISPR/Cas9 system has led to its use in a variety of applications, such as generation of transgenic animals and cell lines and pooled genome-wide screening (27–29)

Recently, CRISPR/Cas9 has been repurposed as a programmable restriction enzyme to direct cleavage in a more precise and customized manner than conventional restriction enzymes, allowing for innovations in cloning and sequencing of complex repeat regions (30–32). Methods using CRISPR/Cas9 as a restriction enzyme have been used to selectively deplete overabundant sequences in a process termed Depletion of Abundant Sequences by Hybridization (DASH) using CRISPR/Cas9 (33). DASH was used to remove targets such as ribosomal RNA (rRNA) from mRNA-seq and wild-type KRAS background sequence from cancer samples by directing their targeted cleavage and preventing their further amplification and sequencing. Similarly, other groups have applied this process to other assays such as the reduction of mitochondrial DNA from ATAC-seq libraries via CARM (CRISPR-Assisted Removal of Mitochondrial DNA) (34, 35). Here, we have adapted and modified the DASH method to deplete adapter dimer and highly-abundant miRNAs from smRNA-seq libraries in a process we have termed miRNA and Adapter Dimer - Depletion of Abundant Sequences by Hybridization (MAD-DASH). MAD-DASH effectively removes these sequences either alone or in combination, increasing proportional read depth of non-targeted species and dramatically improving library construction throughput by enabling SPRI bead size-selection instead of denaturing gels. We identify improvements in the rational design of adapter sequences governing MAD-DASH efficacy and demonstrate the utility of this method for blood based, low-input smRNA-seq biomarker studies with the removal of adapter dimer and a known highly abundant erythrocyte contaminant miRNA from human plasma.

## METHODS

### Total RNA Isolation from Plasma

Peripheral blood sample collections for the isolation of RNA from human plasma were performed as previously described (20) in accordance with the Institutional Review Board at the University of Alabama at Birmingham. Briefly, each collection 5 mL of blood was drawn, centrifuged to isolate plasma and then stored at −80°C until further use. Total RNA was isolated from 1 mL thawed plasma using the Plasma/Serum Circulating and Exosomal RNA Purification Kit (Slurry Format) (Norgen Biotek) and concentrated to 20 uL using the RNA Clean-Up and Concentration Kit (Norgen Biotek). To limit sample variation between MAD-DASH replicate groups, plasma RNA from multiple donors was combined prior to library construction.

### Cas9 and MAD-DASH sgRNA preparation

*S. pyogenes* Cas9 (5 ug/uL, 30 uM) was purchased from PNABio (CPO2). sgRNAs were designed as described in the main text with full sequences listed in Table S1. sgRNAs were constructed following previously described methods (36). Briefly, T7 promoter containing single stranded oligos (Integrated DNA Technologies) corresponding to each sgRNA were annealed to a consistent tracrRNA sequence to generate a double-stranded sgRNA DNA template of the form [TAATACGACTCACTATAGG-N20-GTTTTAGAGCTAGAAATAGCAAGTT-AAAATAAGGCTAGTCCGTTATCAACTTGAAAAAGTGGCACCGAGTCGGTGCTTTT] wherein the N20 is the sgRNA sequence. sgRNA length and variable 5’ G inclusion for efficient transcription was varied as needed for specific sgRNAs. This template was then used for *in vitro* transcription (IVT) using the MEGAshortscript T7 Transcription Kit (Invitrogen). IVT was carried out with 1 ug template at 37°C for 16 hrs, treated with 1 uL TURBO DNASE (Invitrogen) for 15 min, heat inactivated at 95°C for 10 min, and the transcribed RNA was cleaned up using the MEGAclear Transcription Clean-up Kit (Invitrogen). sgRNAs were quanitified using Broad Range RNA Qubit (Invitrogen), normalized to 1.5 ug/uL (46.5 uM) and stored in single-use aliquots at −80°^−^C. Multiplex 5’ adapter dimer targeting sgRNAs were pooled evenly before freezing.

### MAD-DASH Small RNA Sequencing

Small RNA sequencing was performed as described previously (20), with modifications to incorporate the MAD-DASH protocol. A full, detailed MAD-DASH smRNA-seq protocol including sgRNA preparation is in the Supplemental Methods. Oligos for MAD-DASH smRNA-seq were from Integrated DNA Technologies and are listed in Table S1. Briefly, 4 uL isolated total RNA from human plasma or 50 ng purchased Human Brain Total RNA (Invitrogen) was used as input in a ligation reaction (25°C for 1 hr) with 10 uM pre-adenylated 3’ adapter ligation reaction and 1 uL T4 RNA Ligase2, truncated (NEB). This ligation product was then annealed with the 1 uL 10 uM RT primer at 25°C for 1 hr. 5’ adapter ligation was performed at 25°C for 1 hr using 1 uL T4 RNA Ligase 1 (NEB) and 1uL 20 uM multiplex 5’ adapter pool or the modified 5’ adapter (predenatured at 70°C for 2 min). Reverse transcription of ligation products was performed using Superscript II Reverse Transcriptase (Invitrogen) before performing 4 cycles of PCR (PCR1) using Phusion 2X High-Fidelity PCR Master Mix (NEB) with cycling conditions of 94°C for 30s, 4 cycles of 94°C for 10s and 72°C for 45s, and a final extension at 65°C for five minutes. PCR products (50 uL total) were cleaned and concentrated to 10 uL using Agencourt AMPure XP SPRI Beads (Beckman-Coulter).

To perform the adapter dimer or hsa-miR-16-5p MAD-DASH, we followed the instructions from PNABio with modification for incorporation into our smRNA-seq workflow. For a MAD-DASH reaction volume of 20 uL, 1 uL 30 uM Cas9 was pre-incubated with 5 uL 46.5 uM of sgRNA, 2 uL 10X NEBuffer3.1 (NEB) and 2 uL 10X BSA (NEB) at 37°C for 15 min before combining with the 10 uL sample DNA and incubating at 37°C for 2 hrs. After two hours, 1 uL 4 ug/uL RNAseA (NEB) was added to remove sgRNA (37°C for 15 min) followed by 1 uL 20 ug/uL Proteinase K (NEB) (37°C for 15 min, 95°C for 15 min) to rapidly inactivate Cas9. We found this incubation with Proteinase K to be critical for library yield, as Cas9 not only has extremely high DNA-binding when not complexed to an sgRNA, but also as described in the main text there was significant non-target competitive-binding when using adapter dimer targeting sgRNAs. Other methods of Cas9 inactivation including heat (65°C or 95°C) or STOP Solution (30% glycerol, 1% SDS, 250mM EDTA, pH 8.0) were less effective or resulted in significant reductions in final library yield (Figure S1). Proteinase K inactivation at 95°C was critical, as the post-Proteinase K treated samples were immediately used to perform a second round of PCR amplification (PCR2) for 11 cycles using identical cycling conditions. Proteinase K treatment and inactivation rendered post-MAD-DASH sample cleanup prior to PCR2 unnecessary. Following PCR2, samples were once again cleaned up with AMPure XP beads. We term samples cleaned up with 0.9X and 1.8X AMPure bead volumes serially followed by eluting in 20 uL dH20 as “MAD-DASH smRNA-seq protocol samples”. For comparison with no-Cas9/sgRNA MAD-DASH control gel extracted samples, a single sided AMPure bead cleanup using 1.8X was performed eluting in 17 uL. Denaturing gel electrophoresis with 10% TBE-Urea Mini-PROTEAN gels (Bio-Rad) and extraction were performed as described previously, with the only difference being we extracted the ~75 bp region between the ~125 bp adapter dimer band and 200bp band (rRNA) as opposed to just the 145 bp miRNA-library band. To account for this larger gel slice, we doubled our gel soaking solution (2 mL 5M Ammonium Acetate,2 mL 1% SDS solution, 4 uL 0.5M EDTA, 16 mL RNAse/DNAse free dH20) step (2hr at 70°C) and subsequent addition of 100% isopropanol volumes to 600 uL to precipitate the DNA overnight. DNA was washed with ice-cold 80% ethanol before resuspending in 10 uL dH20.

Libraries were quantified with the KAPA Library Quantification Kit for Illumina (KAPA Biosystems) and normalized to 5 nM concentration. MAD-DASH smRNA-seq libraries were combined (2 uL for adapter dimer MAD-DASH samples and 1 uL of samples without adapter dimer depletion to prevent adding proportionally too much adapter dimer to the flow cell) to yield a final 5 nM total pool. MAD-DASH smRNA-seq library pools from (1) AMPure bead cleaned-up brain RNA samples (2) gel-extracted brain RNA samples, or (3) gel-extracted plasma samples were sequenced on 2 lanes each on an Illumina HiSeq2500 with single-end 50bp reads according to standard Illumina protocols.

### Data Analysis

Data analysis was performed as previously described (20) with minor modifications to incorporate MAD-DASH specific details. FASTQs were demultiplexed using custom index sequences and adapter sequences were trimmed using Cutadapt (37). Sequences <15 bp after adapter trimming were separated into a “Short Fail” FASTQ, from which adapter dimer reads (corresponding to a blank sequence line, i.e. trimmed read length of 0 bp) and non-adapter dimer short fail reads were collected. Trimmed reads were aligned to pre-miRNA sequences (miRBase v19 (38)) using Bowtie2 (39) with only two mismatches allowed and keeping only unique best alignments. Mature miRNA counts were determined by counting the aligned pre-miRNA reads overlapping mature mIRNA boundaries using BEDTools (40). Trimmed reads that did not align to miRNAs contained other smRNA sequences and were designated as “non-miRNA usable reads” for purposes of read fraction calculations. Statistical analysis was performed using the R statistical software package (version 3.4.0). For plotting comparisons between groups, replicate libraries were downsampled to equivalent counts and summed before downsampling again with the compared group to yield an equivalent number of aligned reads between groups and then transforming to counts-per-million. Differential expression between samples was compared using the above downsampling process without summing group replicates prior to between group downsampling and using DESeq2 (41) with local dispersion estimates and LRT tests. Significant differential expression was defined as a Benjamini-Hochberg adjusted p-value (FDR) <0.05.

### Adapter Dimer quantitative PCR in human plasma MAD-DASH libraries

qPCR primers were designed to specifically amplify adapter dimer sequences from either multiplex 5’ adapter 1 or the modified 5’ adapter and so they would be incapable of amplifying cleaved adapter dimer sequences (Figure S2). qPCR was performed with Power SYBR Green Master Mix (Invitrogen) in 10 uL reactions using post-PCR2 samples with standard curve using synthetic adapter dimer library to accurately quantify adapter dimer concentration. Multiplex adapter dimer amounts were multiplied by 4 to account for the other 3 adapter dimer sequences resulting from the equimolar multiplex 5’ adapters. Data analysis was performed using R and comparisons between treated and untreated replicate adapter dimer amounts were performed using an unpaired two-sided Wilcox test with significance set as p<0.05.

## RESULTS

### Overview of MAD-DASH

To overcome the experimental limitations and low-throughput of our standard smRNA-seq workflow (20) caused by excessive adapter dimer and overabundant miRNAs, we sought to adapt the DASH procedure to smRNA-seq (Figure 1A). However, technical differences in smRNA-seq library preparation compared to that of RNA-seq used in DASH required several important modifications to the DASH procedure. First, because the Cas9 enzyme almost exclusively depends on double-stranded DNA (dsDNA) for efficient nuclease activity (42), DASH performs CRISPR/Cas9 *in vitro* digestion after mRNA cDNA second-strand synthesis before library amplification. However, in most smRNA-seq workflows, smRNA cDNA generated in a first-strand synthesis is immediately amplified in PCR to generate dsDNA libraries. Because Cas9 is a single-turnover enzyme (43), it is critical that *in vitro* digestion occurs before full amplification to ensure sufficient reduction of target sequences. Therefore, to generate dsDNA libraries amenable to CRISPR/Cas9 targeting, we performed a first PCR with a limited number of cycles (four) before performing MAD-DASH and then further amplifying with a second round of PCR (eleven cycles). Given the single-turnover nature of Cas9 enzymatic activity, this approach ensures the presence of dsDNA required for Cas9 *in vitro* digestion while also maintaining a low DNA-to-Cas9 ratio necessary for efficient targeted sequence removal.

**Figure 1.**
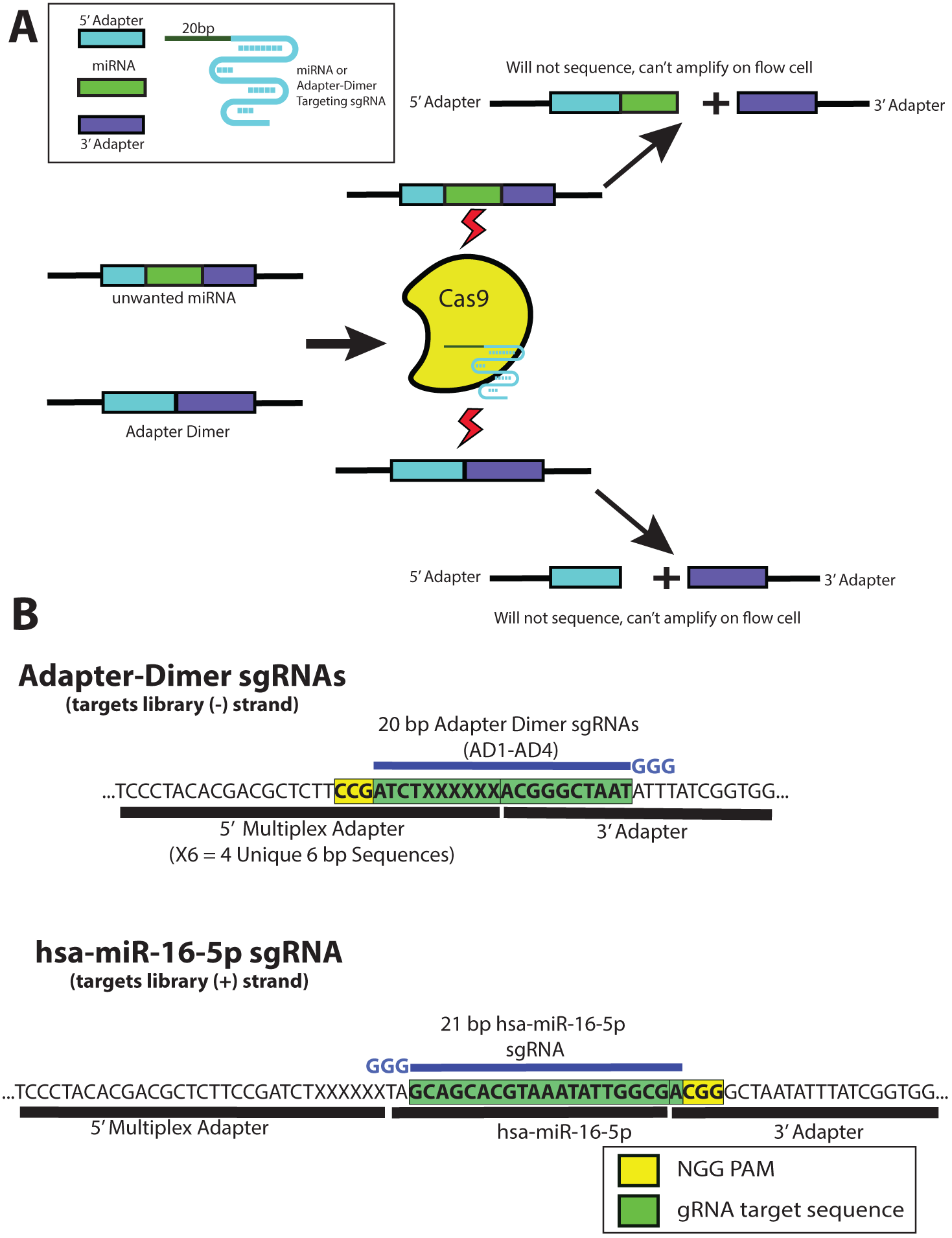
Depletion of adapter dimer and overabundant miRNAs with MAD-DASH. **(A)** MAD-DASH employs Cas9 and adapter dimer- or miRNA-specific sgRNAs to selectively deplete these sequences from final libraries. Cleaved sequences will not amplify during the second round of PCR amplification, and are also no longer suitable substrates for bridge amplification during Illumina sequencing (though they will still be able to bind to the flow-cell using the P5/P7 sequences remaining on each cleaved library). **(B)** Design of adapter dimer and hsa-miR-16-5p sgRNAs using available pre-existing PAM sites on our smRNA-seq 5’- and 3’-adapters. Adapter dimer targeting sgRNAs use a minus strand “NGG” and have ten base pairs of homology to both the 5’- and 3’ adapter, ensuring target cleavage specificity for adapter dimer sequences. hsa-miR-16-5p uses a 3’ PAM site one base away from the 3’ -adapter ligation junction, which can be generalized to other smRNAs and provides highly-specific targeting while minimizing off-targets. Shown are only the plus strand of non-PCR tailed sequences. Green boxes indicate the plus strand sequence corresponding to sgRNA location in the dsDNA library, while the yellow box indicates the plus strand location of the “NGG” PAM in the dsDNA library. sgRNA location is depicted in blue along with the three 5’ “G” nucleotides necessary for high levels of T7 *in vitro* transcription.

Another issue we considered in developing MAD-DASH is the considerably restricted targetable sequence space. Unlike targeting ribosomal RNA derived cDNA or mitochondrial DNA with hundreds of possible PAM sites, smRNA-seq adapters are usually ~25-30 bp in length and the location of PAM sites and possible sgRNAs is limited. Critically, successful targeting of adapter dimer sequences must incorporate sufficient base pair sequence from both the 5’ adapter and the 3’ adapter to prevent off-target cleavage of the rest of the library, which has both a 5’ and 3’ adapter ligated to it. In our smRNA-seq workflow, we use an equimolar pool of four 5’ adapters with a consistent region and a unique 3’ 6 bp base-diverse region to improve smRNA ligation efficiency (predicated on improved adapter-RNA hybridization and resulting favorable ligation cofold structure (16)). To deplete adapter dimer sequences, we designed four unique sgRNAs targeting each possible adapter dimer using a “CGG” PAM site found in the consistent region of the 5’ adapters’ minus strand (Figure 1B). We reasoned that these sgRNAs have a balanced 10 bp of similarity to both their respective 5’ and 3’ adapter sequences and should thus limit off-target cleavage to smRNAs with significant 5’ sequence similarity to the 5’ end of the 3’ adapter. To confirm our sgRNA designs selectively and effectively deplete adapter dimer sequences, we performed Cas9 *in vitro* digestion of synthetic adapter dimer sequences prior to implementation in the full MAD-DASH protocol (Figure S3).

A significant benefit of MAD-DASH relative to other adapter dimer depletion methods is the ability to also deplete specific smRNAs from the library. Similar to depletion of adapter dimer, removal of highly abundant miRNAs such as hsa-miR-16-5p (44) dramatically improves detection of more lowly expressed smRNA species. Utilizing the “CGG” PAM site at the 5’ end of our 3’ adapter allows for highly specific targeting of smRNAs using their 3’-most ~20 bp sequence (Figure 1B) with minimal dependence on adapter nucleotides. Importantly for miRNA depletion, Cas9 nuclease efficiency is typically more sensitive to mismatches in the 8-12 bp PAM-proximal seed region (45). Thus, use of the 3’ adapter PAM site should improve discrimination between miRNA family members that share the same PAM-distal 5’ miRNA seed region. One potential limitation of this 3’ adapter PAM choice is the presence of target miRNA 3’ isomiRs that have 3’ end miRNA sequence variation that can occur due to stochastic DICER pre-miRNA processing or 3’ non-templated base addition and would lead to deleterious PAM-proximal sgRNA-target mismatches (46, 47). Although this phenomenon is not necessarily a concern for all miRNAs, we devised a modified MAD-DASH sgRNA targeting strategy that shows similar efficacy to the 3’ adapter PAM sgRNAs on synthetic miRNA libraries using truncated sgRNAs (48) and amenable miRNA internal PAM sites (present in ~26% of human mature miRNAs annotated in miRBase v21) (Fig S4). However, we have found hsa-miR-165p does not contain significant isomiRs in human plasma RNA, and thus all further hsa-miR-16-5p targeting MAD-DASH performed in this study utilized the above 3’ PAM adapter strategy.

Finally, we considered the dynamics of the inverse relationship between RNA input amount and adapter dimer formation and its effect on MAD-DASH design. In general, low RNA inputs lead to greater adapter dimer formation, which can be especially pronounced in the ~1-10 ng/uL range of RNA from many common clinical biofluids (often comprising > 90% of the amplified library in our experience). Frequently, the dilution of adapters prior to ligation reactions is employed as a means of reducing adapter dimer formation, even when using chemically modified adapters. However, we find that high molar excess of both 3’ and 5’ adapters (10 uM and 20 uM, respectively) is necessary to drive ligation reaction efficiency and more accurate quantification (Figure S5). Conversely, there is a direct relationship to RNA input and targeted miRNA species. Therefore, both high- and low-RNA inputs will face specific challenges regarding sufficient removal of abundant miRNAs or adapter dimer, respectively. To demonstrate the effectiveness of adapter dimer- and miRNA-targeting MAD-DASH in the low RNA-input range used in many clinical sequencing projects, we have constructed smRNA-seq libraries from total brain RNA (50 ng) and human plasma RNA (~1-10 ng), which yielded ~2 nM and ~0.2 nM of adapter ligated product prior to MAD-DASH *in vitro* digestion. For purposes of determining amounts of excess Cas9 and sgRNA relative to target DNA when targeting adapter dimer, we treated the library as if it contained 100% adapter dimer. While this is certainly an overestimation of true adapter dimer concentration, we reasoned that because Cas9/sgRNA DNA-binding activity is more permissive to sequence variation and distinct from its nuclease activity (43, 49, 50), the presence of significant sgRNA sequence similarity on all adapter-ligated sequences would yield considerable competitive off-target Cas9/sgRNA binding (but not cleavage). We thus used a high molar excess of Cas9 input (5 ug, 30 uM) and adapter dimer targeting sgRNA input (7.5 ug, 232.5 uM), which yielded final molar excess relative to target of ~1,500X / ~6,000X and ~15,000X / ~60,000X in brain and plasma RNA, respectively. Identical Cas9/sgRNA input amounts were used when targeting hsa-miR-16-5p (likely considerably more in excess to target), while multiplexing of adapter dimer and hsa-miR-16-5p targeting sgRNA used an identical Cas9 amount and equal amounts (20%) of each sgRNA in the pool. These ratios represent a roughly 10-fold increase of those used in DASH and other sequencing library CRISPR/Cas9 *in vitro* digestion methods. However, we found this increase to render the MAD-DASH protocol robust when depleting large amounts of adapter dimer and miRNAs and simplify Cas9/sgRNA-to-target ratios. We expect that varying RNA input during smRNA-seq library construction may require more or potentially less Cas9/sgRNA in the *in vitro* digestion.

Successful depletion of adapter dimer from smRNA-seq libraries obviates the need to perform low-throughput denaturing gel separation and cleanup, and consequently our MAD-DASH smRNA-seq protocol employs a double-sided SPRI bead size selection for library sequences less than ~200 bp, corresponding to insert range up to approximately 75 bp. Because all library preparation steps including MAD-DASH and final library cleanup can be done in a 96-well plate using SPRI beads, throughput is dramatically increased and amenable to automation. We estimate that average library construction time for a full plate of 96 samples (requiring 6 denaturing gels followed by extraction and precipitation in our standard protocol) could be completed in one day for MAD-DASH smRNA-seq compared to 3-4 days with our standard protocol.

### Evaluating the quantitative performance of MAD-DASH in brain total RNA

To evaluate the effect of reducing adapter dimer and a representative highly-abundant miRNA (hsa-miR-16-5p) both alone and in combination, we generated replicate MAD-DASH no-Cas9/sgRNA control, adapter dimer MAD-DASH, hsa-miR-16-5p MAD-DASH and combined adapter dimer / hsa-miR-16-5p MAD-DASH smRNA-seq libraries and analyzed the normalized abundances of specific sample fractions (Figure 2A) as determined by DESeq2 (see Materials and Methods). MAD-DASH significantly depleted adapter dimer (6.5X) and hsa mir-16-5p (52.1X) alone and in combination (7.5X/6.5X), while increasing non-hsa-miR-16-5p miRNA and non-miRNA usable reads as much as 23%. MAD-DASH targeted adapter dimer sequences were depleted to less than 1.6% of total read fraction, while hsa-miR-16-5p was depleted to as little as 0.015% of total read fraction. We limited further in-depth differential expression analysis to changes in miRNAs and adapter dimer sequences because alignment of non-miRNA smRNAs to the full reference assembly can be complicated (51) and was not expected to demonstrate significantly different results.

**Figure 2.**
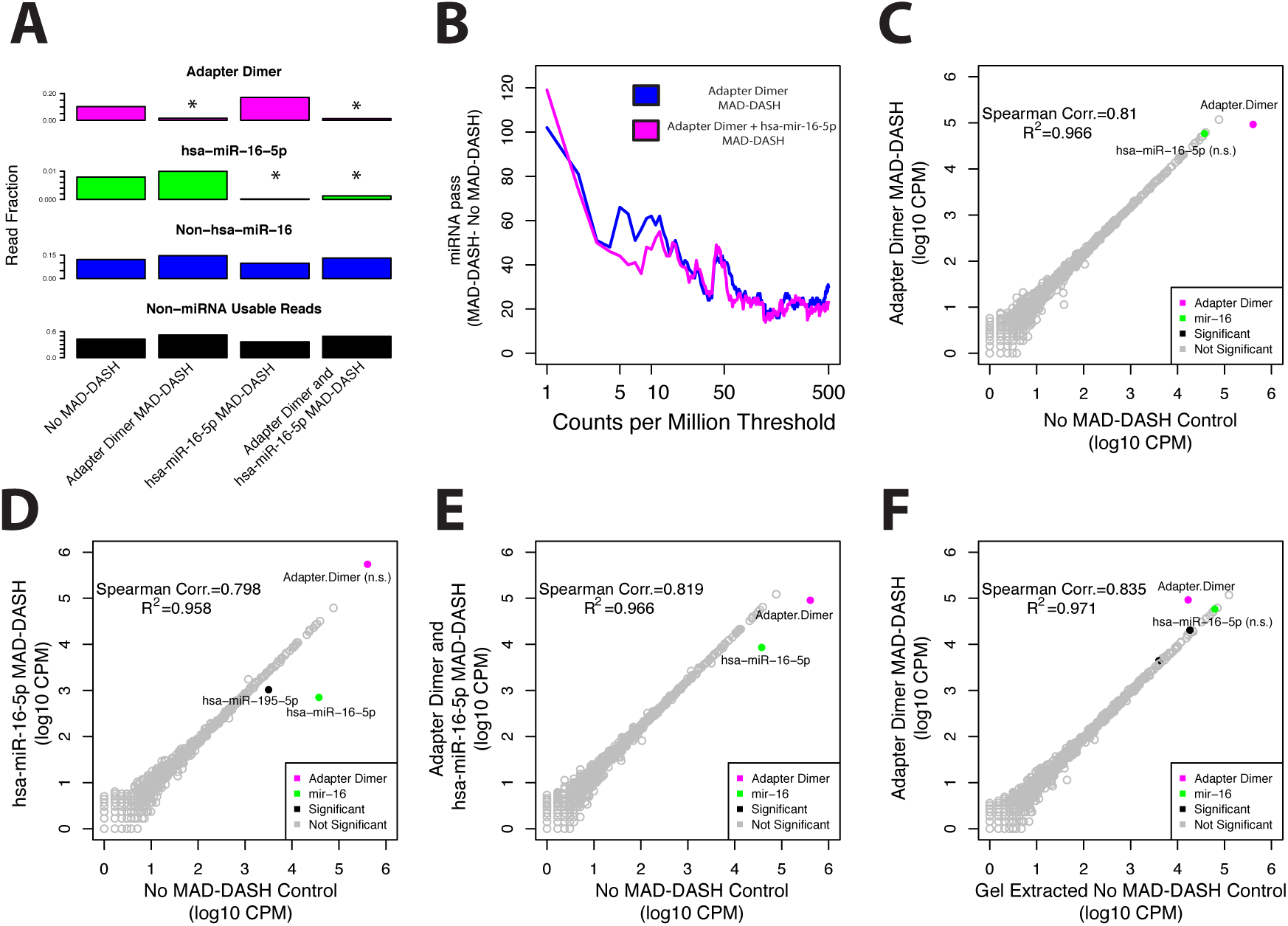
MAD-DASH smRNA-seq reduces adapter dimer and hsa-miR-16-5p in human brain total RNA samples. **(A)** Normalized read fraction of downsampled and CPM normalized adapter dimer sequences, hsa-miR-16-5p, non-hsa-miR-16-5p miRNAs, and non-miRNA usable reads (see Materials and Methods). MAD-DASH significantly depletes adapter dimer (6.5X) and hsa miR-16-5p (52.1X) when targeted individually and 7.5X/6.5X when targeted simultaneously. For adapter dimer or hsa-miR-16-5p, asterisks indicate that a given read fraction was determined to be significantly altered between groups using DESeq2 (FDR <0.05). **(B**) Plot depicting the difference in the number of mature miRNA species reaching a specified count per million threshold between adapter dimer MAD-DASH smRNA-seq replicates and no-Cas9/sgRNA MAD-DASH control replicates. Downsampling and CPM normalization was performed as described in Materials and Methods. **(C-F)** Read counts from normalized replicate groups for treated vs control MAD-DASH samples targeting (C) adapter dimer (D) hsa-miR-16-5p and (E) adapter dimer and hsa-miR-16-5p (F) adapter dimer compared to gel extracted no-Cas9/sgRNA control. Adapter dimer is shown in magenta and hsa-miR-16-5p is shown in green, and their counts are significantly different between samples unless indicate with “(n.s)”, e.g. non-significant. Other significantly different miRNAs are colored black, with those having a log_2_-fold-change > 1 being labeled with text. Non-significantly different miRNAs are depicted as open gray circles. Significance was determined with DESeq2 and set as a Benjamini-Hochberg corrected p-value < 0.05.

Like the effect observed when using our 5’ hairpin blocking method (20), depletion of highly abundant adapter dimer and hsa-miR-16-5p results in increased sensitivity for lowly abundant species (Figure 2B). We detected 102 more miRNAs compared to untreated replicates, with as many as 62 more at the commonly used threshold of ten counts per million (CPM). Comparison of significant differential expression between MAD-DASH treated and control replicates demonstrated highly-specific depletion of targeted sequences (Figure 2C-2E). No significant off-targets were observed when using adapter dimer sgRNAs while hsa-miR-16-5p sgRNA use resulted in only one significant off target, hsa-mir-195-5p. hsa-mir-195-5p is a member of the hsa-mir-16-5p seed region family and shares nearly identical sequence to hsa-mir-16-5p (Table S2). Interestingly, when hybridized to an hsa-mir-195-5p library the hsa-mir-16-5p sgRNA contains a one base pair insertion “bulge” at the PAM -2 site, which appears to be well tolerated and is consistent with prior observations regarding the effect of sgRNA/DNA insertion/deletions on Cas9 nuclease activity (52).

To demonstrate that the MAD-DASH smRNA-seq workflow’s implementation of a SPRI bead cleanup did not negatively affect library reproducibility compared to our standard gel extraction method, we compared the adapter dimer MAD-DASH replicates to gel extracted MAD-DASH no-Cas9/sgRNA control replicates (size extracted 130bp-200bp) (Figure 2F). Only two miRNAs (hsa-miR-22-3p and hsa-miR-9-3p) were significantly altered but had minimal log2 fold-changes (~0.2). While the adapter dimer fraction in the MAD-DASH samples was reduced 6.5-fold to less than 1.6% of the library, it was still roughly 5X more abundant than adapter dimer in the gel extracted samples. Nevertheless, this level of reduction is likely to be more than sufficient to take advantage of the throughput improvements of the MAD-DASH protocol with minimal increase in cost and sequencing depth.

### Rational design of a modified 5’ adapter improves MAD-DASH adapter dimer depletion

Although our first iteration of the MAD-DASH smRNA-seq method successfully depletes adapter dimer, we sought to optimize the reduction further through rationally modifying the design of our 5’ adapter PAM location. As mentioned, our multiplex adapter dimer sgRNA contains ten base pairs of sequence similarity to both the 3’ adapter and its respective 5’ adapter. Prior work has demonstrated that not only does non-sgRNA bound apo-Cas9 possess considerable nonspecific DNA binding ability, but the Cas9/sgRNA complex is capable of binding competing target-DNA with twelve matching PAM-proximal bases as strongly as perfect target sequence (approximately 1000X longer than complete mismatch) (43). Our multiplex adapter dimer sgRNAs, which have a 10 bp competitor mismatch - 10 bp PAM proximal match - PAM design, are thus expected to have considerable, lasting off-target binding to competitor DNA (every non-target molecule in the library) while not having sufficient sequence similarity to engage the Cas9 HNH/RuvC nuclease domains (49).

With this dynamic in mind, we designed a single 5’ adapter identical to one of our four multiplex 5’ adapters, save for a single extra C at the start of the base diverse region that generated a “CCA” plus strand / minus strand “NGG” PAM site 4 bp from the 5’ adapter / 3’ adapter junction (Figure 3A). Our 19bp modified 5’ adapter sgRNA has a 15 bp competitor mismatch - 4 bp PAM proximal match - PAM design, and is thus expected to exhibit considerably greater adapter dimer depletion by limiting competitive binding to all other adapter ligated sequences. We generated replicate modified adapter MAD-DASH no-Cas9/sgRNA control and modified adapter dimer MAD-DASH libraries that as expected demonstrated significantly greater adapter dimer depletion compared to our multiplex 5’ adapter samples (Figure 3B). Although our single modified 5’ adapter generated considerably more adapter dimer (likely due to less efficient capture of miRNAs caused by only one sequence dictating hybridization and adapter/RNA cofold structure), we were able to deplete adapter dimer 20.3X to less than 1% of the final library. This represents a ~3X greater reduction than in multiplex adapter dimer MAD-DASH samples, and leads to detection of as many as ~101 additional miRNAs (29 at 10 CPM) compared to modified adapter control samples (Figure S6).

**Figure 3.**
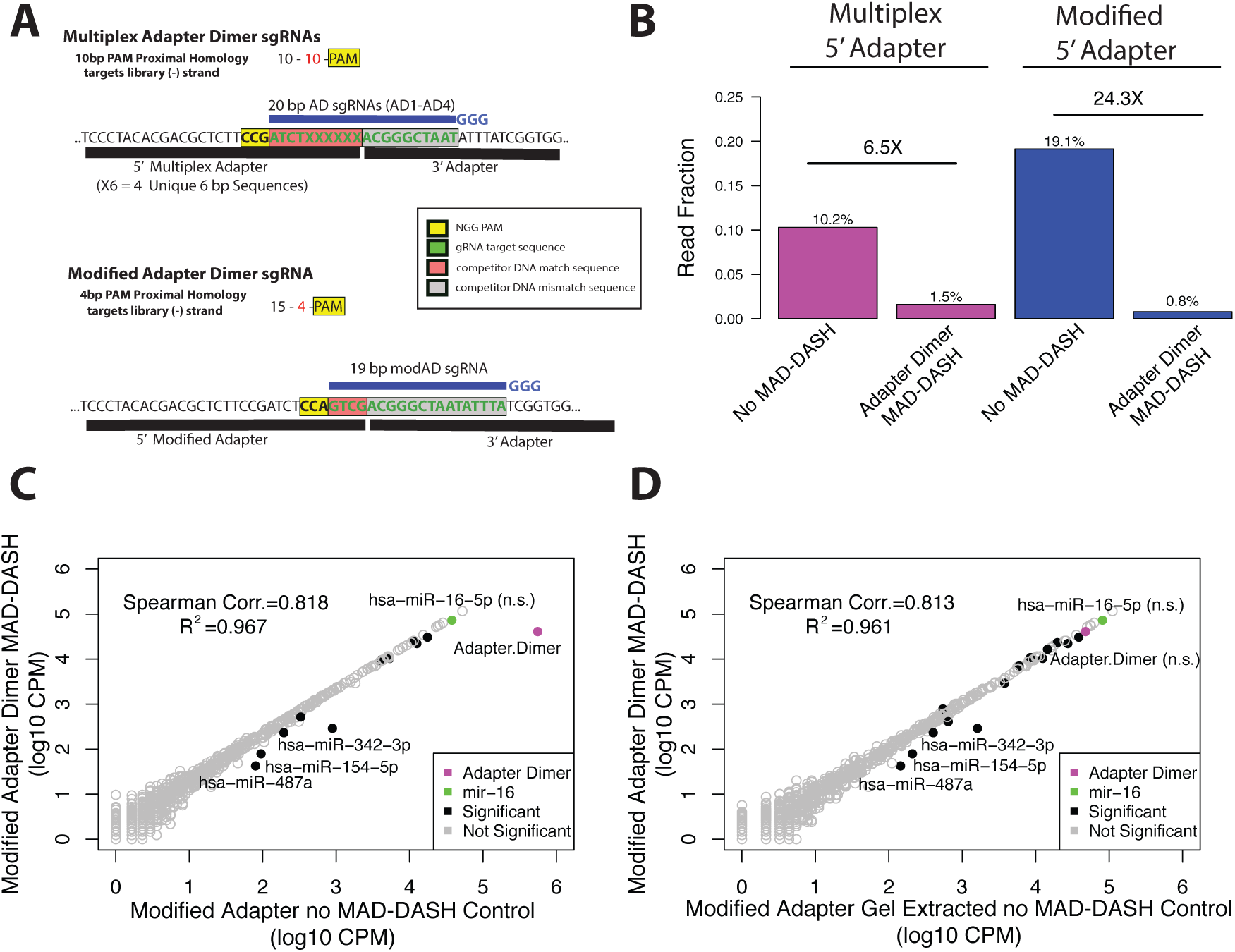
Rational design of a modified 5’ adapter with alternate PAM site enhances depletion of adapter dimer with MAD-DASH. **(A)** Design of the modified 5’ adapter compared to the multiplex 5’ adapter. Compared to the 10 bp competitor mismatch −10 bp competitor match -PAM design used in our first MAD-DASH iteration, the modified adapter uses 15 bp competitor mismatch - 4 bp competitor match-PAM design and is predicted to have as much as 100-fold less binding to other non-target sequences in the library. Shown are the plus strands of non-PCR tailed sequences. Red boxes indicate the plus strand sequence corresponding to competitor match sequence in the dsDNA library, while the grey box indicates the sequence corresponding to sequence that drives adapter dimer target specificity, i.e. competitor mismatch sequence. The yellow box indicates the plus strand location of the “NGG” PAM in the dsDNA library. sgRNA location is depicted in blue along with the three 5’ “G” nucleotides necessary for high levels of T7 *in vitro* transcription. **(B)** MAD-DASH smRNA-seq using the modified 5’ adapter demonstrates substantially greater depletion of adapter dimer, despite making more adapter dimer in the no-Cas9/sgRNA control group. Normalized read fraction of downsampled and CPM normalized adapter dimer sequences for both adapter strategies is depicted. Reductions of adapter dimer in both the multiplex adapter samples and the modified adapter samples were significant (DESeq2 adjusted p-value <0.05). **(C-D)** Read counts from normalized replicate treated vs control groups prepared using the modified 5’ adapter. MAD-DASH samples depleted of (C) modified adapter dimer and (D) modified adapter dimer compared to gel extracted no-Cas9/sgRNA modified adapter control are shown. Adapter dimer is shown in magenta and hsa-miR-16-5p is shown in green. Their counts are significantly different between samples unless indicated with “(n.s)”. Other significantly different miRNAs are colored black, with those having a log_2_-fold-change > 1 labeled with text. Nonsignificantly different miRNAs are depicted as open gray circles. Significance was determined with DESeq2 and set as a Benjamini-Hochberg corrected p-value < 0.05.

Again, comparison of significant differential expression between modified adapter dimer MAD-DASH treated and control replicates demonstrated highly specific depletion, albeit with more off-targets than our multiplex adapter MAD-DASH samples (Figure 3C). This is to be expected due to the reduced number of PAM-proximal bases corresponding to the modified adapter’s minus strand “CAGC” needed for off target mismatch (6 bp vs 12 bp, i.e. NGG + 4 PAM proximal bases or 10 PAM proximal bases). Significantly altered sequences with log2 fold changes greater >1 (hsa-miR-342-3p, hsa-miR-154-5p and hsa-miR-487-3p) all contained “CCN” corresponding to a target strand “NGG” PAM site in the last 7 bp of their sequence (Table S2). Interestingly, hsa-miR-487-3p and hsa-miR-342-3p demonstrate a similar insertion/deletion sgRNA/DNA bulge phenomenon to that seen with hsa-miR-195-5p and our hsa-miR-16-5p targeting sgRNA. Importantly, modified adapter dimer MAD-DASH samples have an equivalent reduction in adapter dimer amounts compared to gel extracted control samples with minimal effect on library representation, achieving the desired technical correspondence between our standard smRNA-seq protocol and MAD-DASH smRNA-seq (Figure 3D) while dramatically increasing library throughput.

### Evaluating MAD-DASH quantitative performance in human plasma RNA

We next sought to demonstrate the utility of MAD-DASH in clinically relevant samples. The detection of circulating smRNAs in human plasma for discovery of diagnostic and prognostic biomarkers is an emerging field. However, as mentioned previously the amount of RNA isolated from patient plasma samples is minimal, on the order of a few ng RNA / mL plasma, leading to low RNA inputs in smRNA-seq and high levels of adapter dimer formation. Additionally, certain blood cell specific miRNAs such as hsa-miR-16-5p and hsa-miR-150 are highly abundant contaminants that limit sensitivity (20, 44). We therefore applied MAD-DASH smRNA-seq using both our multiplex 5’ adapter and our modified 5’ adapter strategy to replicate libraries generated from human plasma samples to demonstrate the successful removal of adapter dimer and hsa-mir16-5p and increase detection of lowly abundant species.

Because no-Cas9/sgRNA MAD-DASH control plasma samples yield >95% adapter dimer, dominating the sequencing reads making interpretation impossible, we instead designed a qPCR assay (Figure S3) to quantify the reduction in adapter dimer between MAD-DASH control and treated samples (Figure 4A). We observed significant levels of proportional reduction when using either the multiplex 5’ adapter (Wilcox P-value < 1.1×10^−3^) or modified 5’ adapter strategy (Wilcox P-value < 7.4×10^−7^), with the modified 5’ adapter MAD-DASH again exhibiting comparatively greater reduction. To confirm specific depletion of highly abundant hsa-miR-16-5p, we sequenced gel-extracted plasma MAD-DASH samples which demonstrated similar specificity to that of brain RNA samples, with only hsa-miR-195-5p as an off-target (Figure 4B). In the case of the modified adapter MAD-DASH samples, there was still a significant reduction in residual adapter dimer remaining after gel extraction (Figure 4C). These reductions in abundant species yield a dramatic increase in detection of possibly informative, lowly expressed miRNAs (Figure 4D) and demonstrate the utility of MAD-DASH for improving the sensitivity of smRNA-seq for lowly-expressed circulating biomarker detection. In principle, extension of MAD-DASH to smRNA-seq samples generated from other biofluids such as cerebrospinal fluid should necessitate only the initial detection of non-informative highly abundant sequences to be depleted along with adapter dimer for greatest effect.

**Figure 4.**
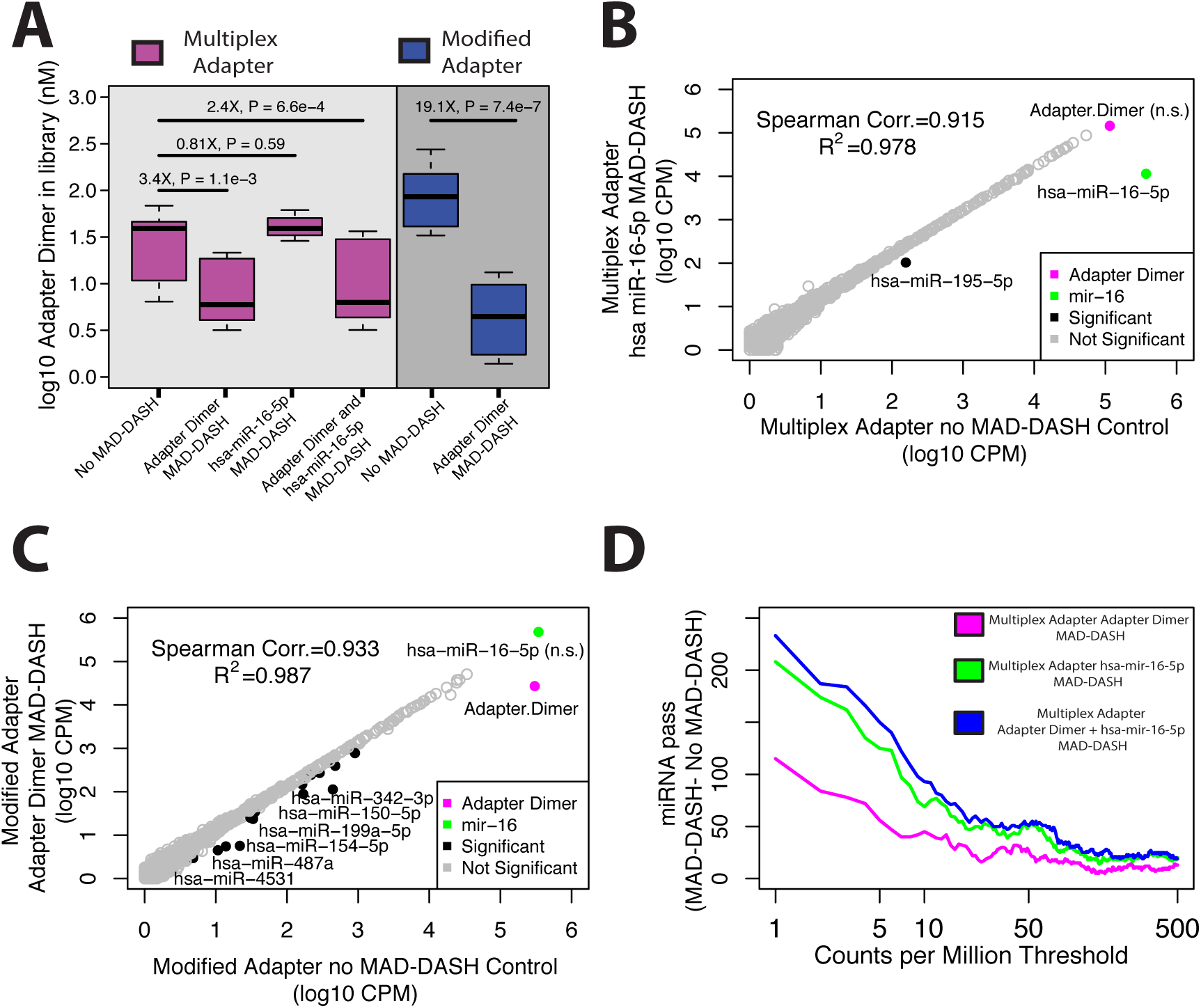
MAD-DASH smRNA-seq reduces adapter dimer and hsa-miR-16-5p in human plasma samples. **(A)** Adapter dimer concentrations in control or treated MAD-DASH smRNA-seq replicates (n=3 in each group) made using the multiplex 5’ adapter (magenta) or the modified 5’ adapter (blue) were determined using qPCR and a synthetic adapter dimer library standard. Multiplex adapter sample concentration were adjusted 4x to account for the three other equimolar multiplex adapters. Similar levels of reduction are seen in MAD-DASH smRNA-seq libraries generated from approximately 5-50 more concentrated brain total RNA samples (see Figure 2). **(B-C)** Sequencing read counts from gel extracted, normalized replicate groups are plotted for treated vs control MAD-DASH samples using either (B) multiplex adapters and targeting hsa-miR-16-5p, or using (C) modified adapter and targeting adapter dimer. hsa-miR-16-5p depletion would not expected to vary between multiplex vs. modified adapter use due to an identical sgRNA / PAM location. Adapter dimer depletion depicted in (C) represents a depletion of residual adapter dimer remaining after gel extraction, indicating that modified adapter dimer MAD-DASH can yield significant library improvements even when using traditional library cleanup methods. **(D)** Plot depicting the difference in the number of mature miRNA species reaching a specified count per million threshold between gel extracted, treated and control MAD-DASH smRNA-seq replicates made using the multiplex adapter and targeting adapter dimer and hsa-mir-16-5p alone and in combination. As in (C), MAD-DASH samples targeting adapter dimer alone and combined with hsa-mir-16-5p depict increased sensitivity even when performing gel extraction. Downsampling and CPM normalization was performed as described in Materials and Methods.

## DISCUSSION

In this study, we have adapted CRISPR/Cas9 *in vitro* digestion for the removal of adapter dimer and highly-abundant smRNA species during smRNA-seq library preparation. These sequences limit the ability to detect lowly-expressed smRNAs and in the case of adapter dimer necessitate arduous, low-throughput denaturing gel size-selection steps. MAD-DASH is a single, simple workflow that renders the need for gel extraction unnecessary in favor of a SPRI bead based size-selection (Figure 5). We estimate that the increased library construction cost when incorporating MAD-DASH is less than $10 per sample using commercially purchased Cas9 and in vitro transcription kits. These costs are likely to be easily covered by the dramatic increase in throughput and ensuing time savings in large scale sequencing projects. This may be particularly true if automation is employed, which MAD-DASH makes possible by allowing for all steps to be performed “on-plate”. We have designed and optimized the MAD-DASH protocol specifically for smRNA-seq, though like the original DASH CRISPR/Cas9 *in vitro* digestion methodology it is inherently generalizable to other library preparation methods with significant adapter dimer amounts. Frequently, however, adapter dimers are much smaller and more easily separable from insert-containing libraries in other applications, underscoring the special utility of MAD-DASH in smRNA-seq.

**Figure 5.**
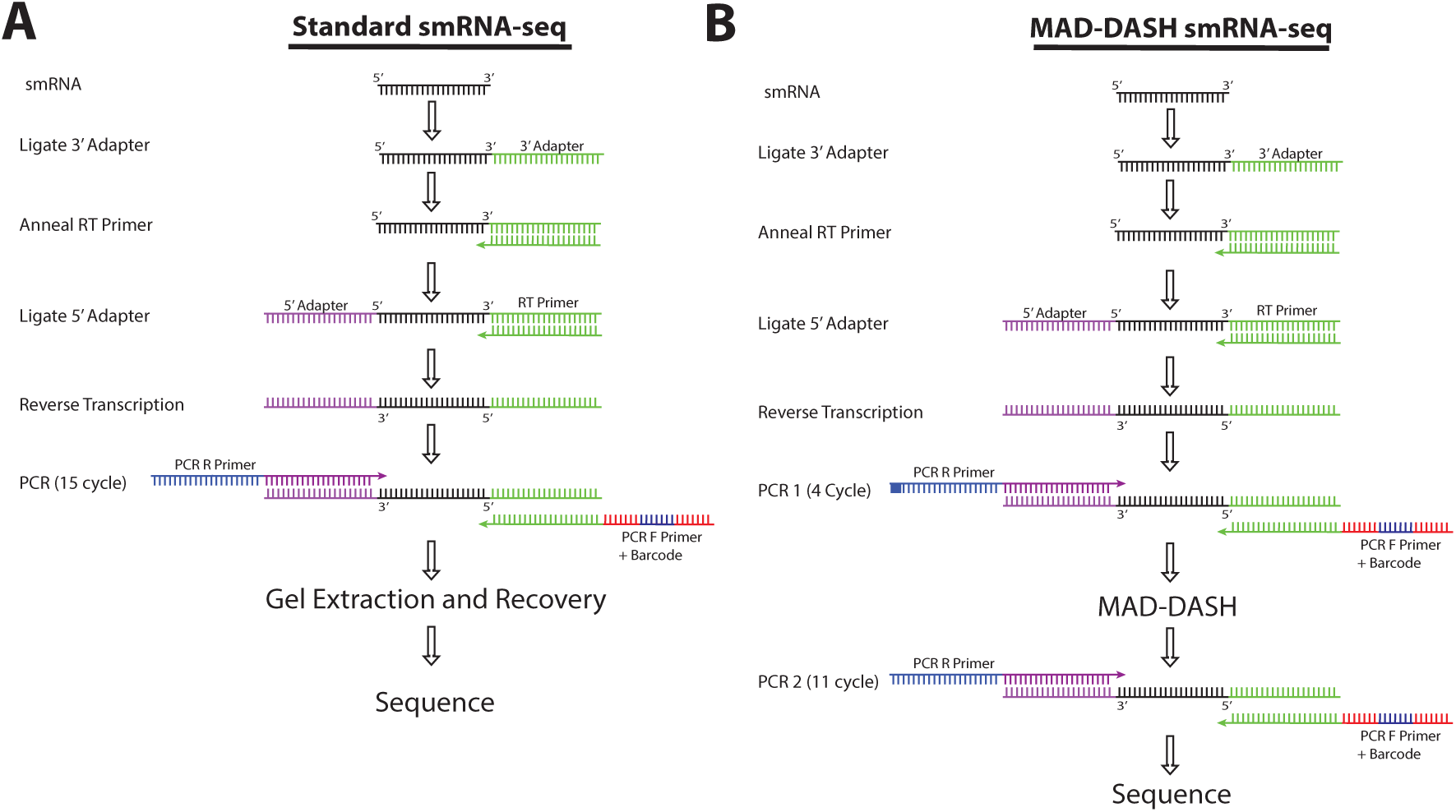
Illustration of standard smRNA-seq (A) and MAD-DASH smRNA-seq (B) workflows. While MAD-DASH involves an extra PCR amplification in addition to the MAD-DASH CRISPR/Cas9 *in vitro* digestion, the extra ~3 hours to accomplish this relative to finishing the PCR in our standard protocol are more than offset by no longer needing to perform gel extraction, DNA recovery and library concentration. For nearly a hundred samples, we estimate that researcher time for smRNA-seq library construction can be reduced to only 1 day using MAD-DASH smRNA-seq and that this throughput can be further improved with greater protocol familiarity or automation.

We demonstrate that our MAD-DASH smRNA-seq library protocol can deplete adapter dimer and targeted miRNAs with very high specificity that is consistent with known sequence-dependent CRISPR/Cas9 on-/off-target discrimination. We show ability to deplete adapter dimer up to 20X to less than 1% of library fraction and reduce abundant hsa-miR-16-5p up to 44X in human plasma RNA to below less than 1%. Additionally, we have demonstrated a rational basis for modifying PAM site and resulting adapter dimer targeting sgRNA location to address the uniquely problematic competitive-binding dynamics present in MAD-DASH smRNA-seq that result from every sequence in the library having significant sequence similarity. While our modified 5’ adapter MAD-DASH exhibits predictably more off-targets, these effects are expected and consistent. While outside the scope of this study, it is expected that further optimizing adapter design with both the dynamics of smRNA ligation efficiency and MAD-DASH depletion efficiency in mind will lead to further improvements in MAD-DASH implementation. Substitution of S. *pyogenes* Cas9 with other Cas9 variants or Cas9 with engineered alternative PAM specificities (53, 54) can expand the limited targeting space in MAD-DASH, allowing for more customizable workflows. Although RNA-targeting CRISPR enzymes have recently been described (55, 56) and could provide a substitute for Cas9 in an RNA-targeting MAD-DASH protocol, these enzymes have been shown to have considerable non-specific RNA cleavage when used *in vitro* which would almost assuredly reduce MAD-DASH specificity and reproducibility. Nevertheless, the most common commercial smRNA-seq library generation kits use “NGG” PAM-site containing adapters that are completely consistent with implementation of MAD-DASH with few changes to their published protocols (Table S3). Importantly, because MAD-DASH can not only remove adapter dimer but also other abundant smRNAs, it can also yield improvements in smRNA-seq methods already designed to limit adapter dimer formation by using chemically-modified adapters or employing template-switching instead of adapter ligation.

Despite its utility, certain limitations in the use of MAD-DASH exist. The impact of the dynamics of competitive-binding on MAD-DASH when using other adapters remains to be explored. Additionally, the use of MAD-DASH with singlecell levels of RNA (in the 10 pg range) is likely to require further optimization. The use of dual randomer sequences in adapters to improve ligation efficiency and thus accurate miRNA representation (16) is likely to hinder the use of MAD-DASH for adapter dimer depletion unless care is taken to design a consistent 20bp sgRNA targetable region. Finally, as described for other CRISPR/Cas9 based depletion methods (33, 35), cleaved sequences still contain intact P5 and P7 flow-cell binding sequences, which although not able to be amplified during bridge-amplification can still bind to the flow cell during initial hybridization. While our cleaved adapter sequences are small enough to be partially removed by the lower limit of SPRI bead binding, MAD-DASH samples may require sequencing a greater library concentration relative to conventional methods. The extent to which these possible technical issues impact library quality and their resolution is the subject of future work. In conclusion, our MAD-DASH smRNA-seq protocol provides a robust, high-throughput method to deplete adapter dimer and unwanted highly-abundant smRNAs in a manner analogous to rRNA and globin reduction from mRNA-seq libraries and thus overcomes a significant obstacle in the use of smRNA-seq in large scale sequencing projects.

## ACKNOWLEDGEMENTS

We thank the patients who have graciously provided plasma samples for this study and members of the Clinical Research Support Program and the Center for Clinical and Translational Science Laboratory at the University of Alabama at Birmingham for their help with sample collection and processing, in particular C. Mel Wilcox, Meredith B. Fitz-Gerald and Robert P. Kimberly. We thank the Genomic Services Laboratory, led by Dr. Shawn Levy, at the HudsonAlpha Institute for Biotechnology for provided sequencing services and data processing. We thank Gregory Cooper and Sara Cooper for critical reading of the manuscript, and also thank Kenneth Day for technical advice with qPCR experiments.

---

**Supplementary Figure S1.**
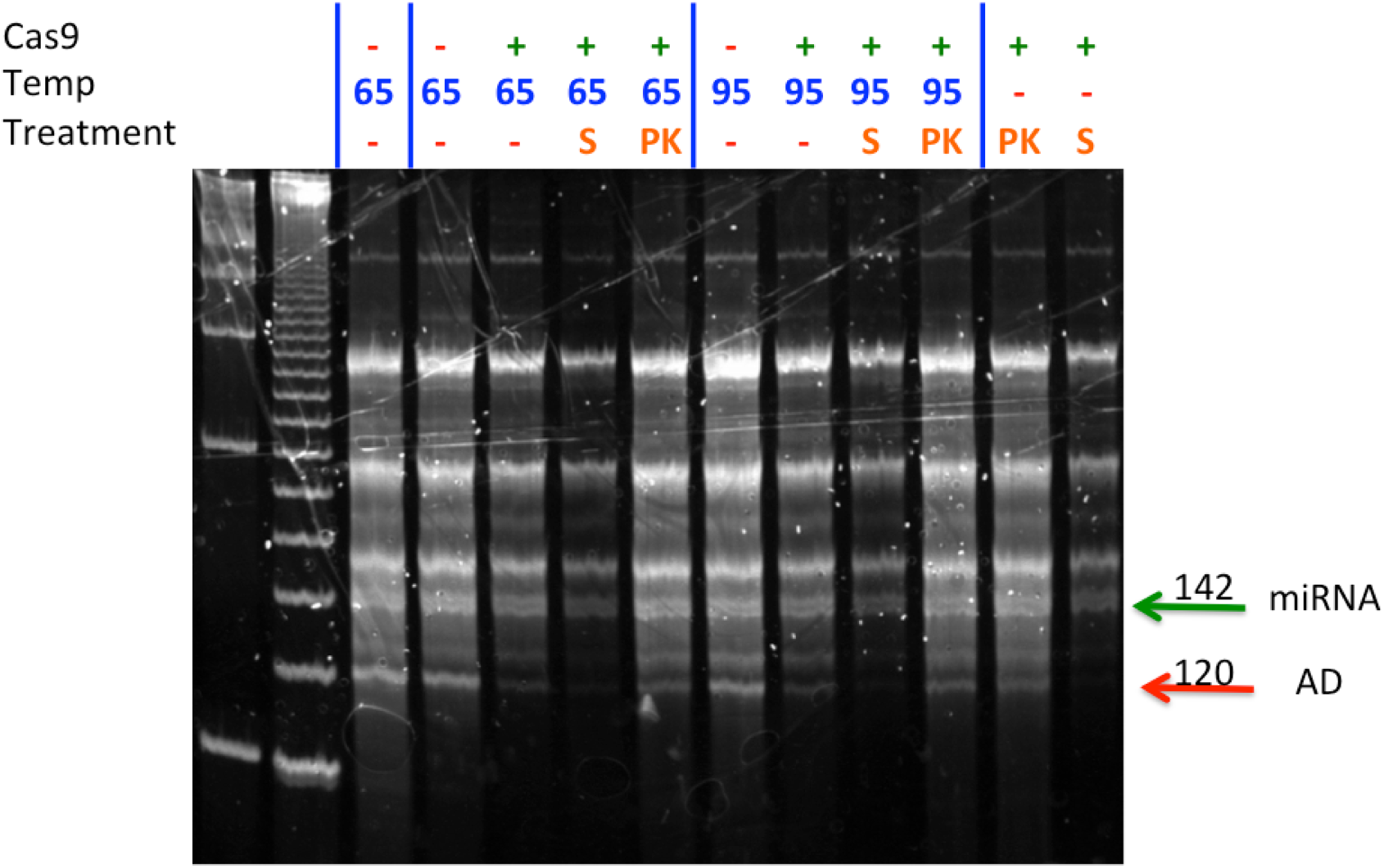
The effect of various experimental conditions to stop the MAD-DASH CRISPR/Cas9 *in vitro* digestion on final library yield. Samples were treated with 1ug Cas9 and either 10 uL multiplex adapter sgRNA and were either not heat inactivated or heat inactivated at 65°C or 95°C, while also varying additional treatments of either 1 uL of 20 ug/uL Proteinase K, 1 uL PNABio STOP Solution. STOP solution greatly impacts final library yield, while heat inactivation alone is insufficient to dissociate the Cas9/sgRNA complex from library DNA, which leads to relatively greater amounts of adapter dimer in the library compared to the combination of heat and proteinase K.

**Supplementary Figure S2.**
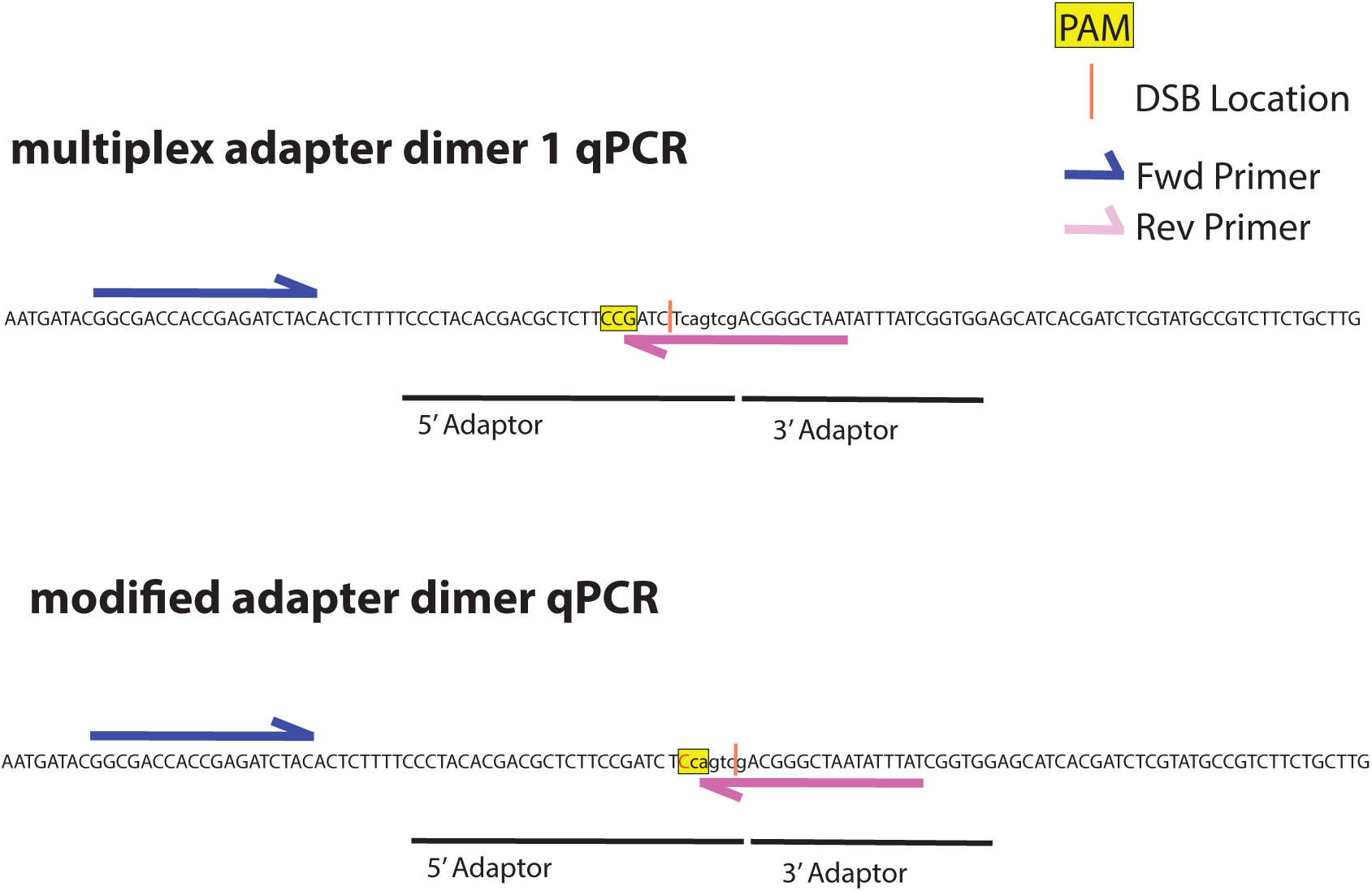
Design of adapter dimer qPCR primers to assess adapter dimer concentrations after PCR2. A consistent forward primer to amplify both multiplex and modified adapter dimers was designed, with adapter specificity coming from the reverse primer. The reverse primers were designed to both incorporate some amount of 5’ adapter and 3’ adapter sequences while also spanning the MAD-DASH CRISPR/Cas9 mediated double-stranded break, such that they had only four base pairs with which to anneal to non-cleaved adapter dimer, which prevents non-specific amplification of non-adapter dimer species. Furthermore, mismatches in the 3’ head of primers are typically result in no or inefficient amplification. Matching Off-target sequences with other sequences in the library would require significant sequence homology and would not be expected to vary between treated and untreated samples.

**Supplementary Figure S3.**
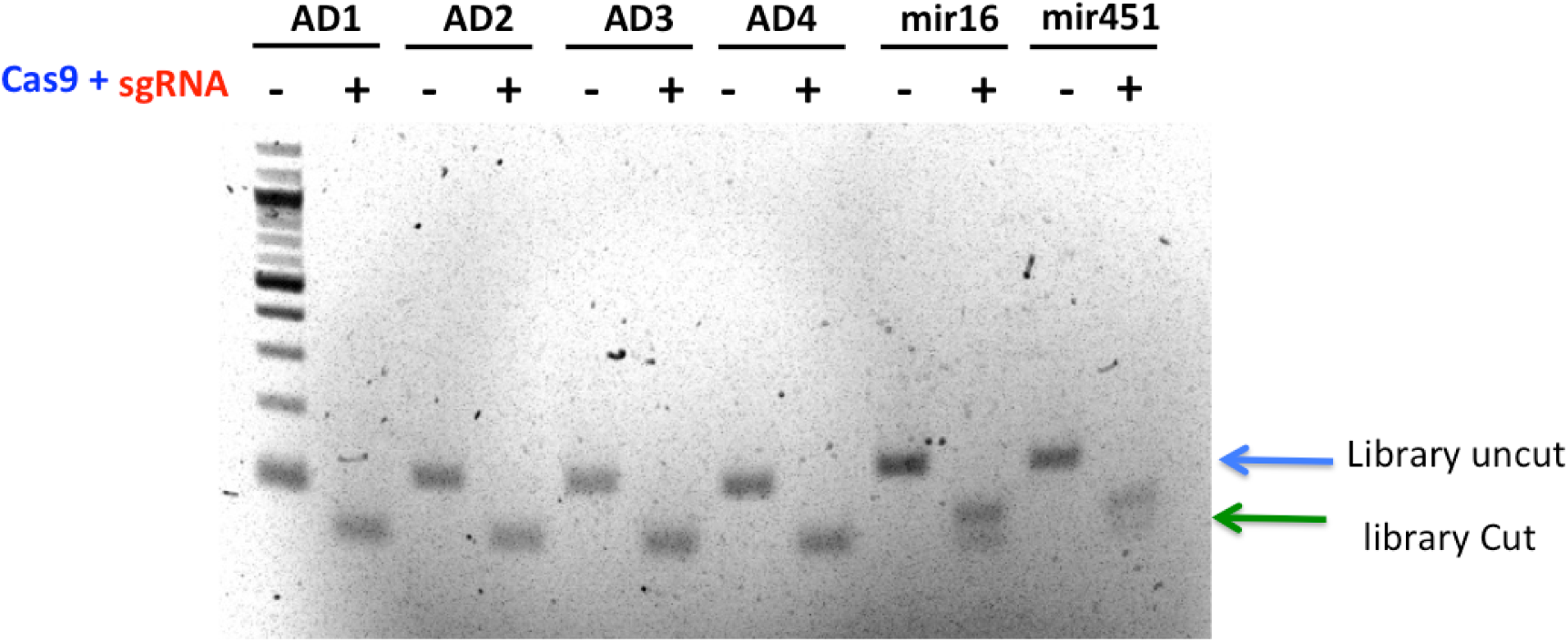
Initial MAD-DASH Cas9/sgRNA *in vitro* digestion assay for all multiplex adapter dimer, hsa-miR-16-5p and hsa-miR-451a gBlocks. Uncleaved and MAD-DASH cleaved libraries are depicted with expected base pair sizes. 1ug Cas9 and 300 ng sgRNA were used for optimization and achieve nearly 100% cleavage in all samples. Due to well number limitations, uncut multiplex adapter dimer 1 MAD-DASH test was run in the 100 bp ladder well, lane 1, but was deemed not important for interpretation of the *in vitro* digestion assay.

**Supplementary Figure S4.**
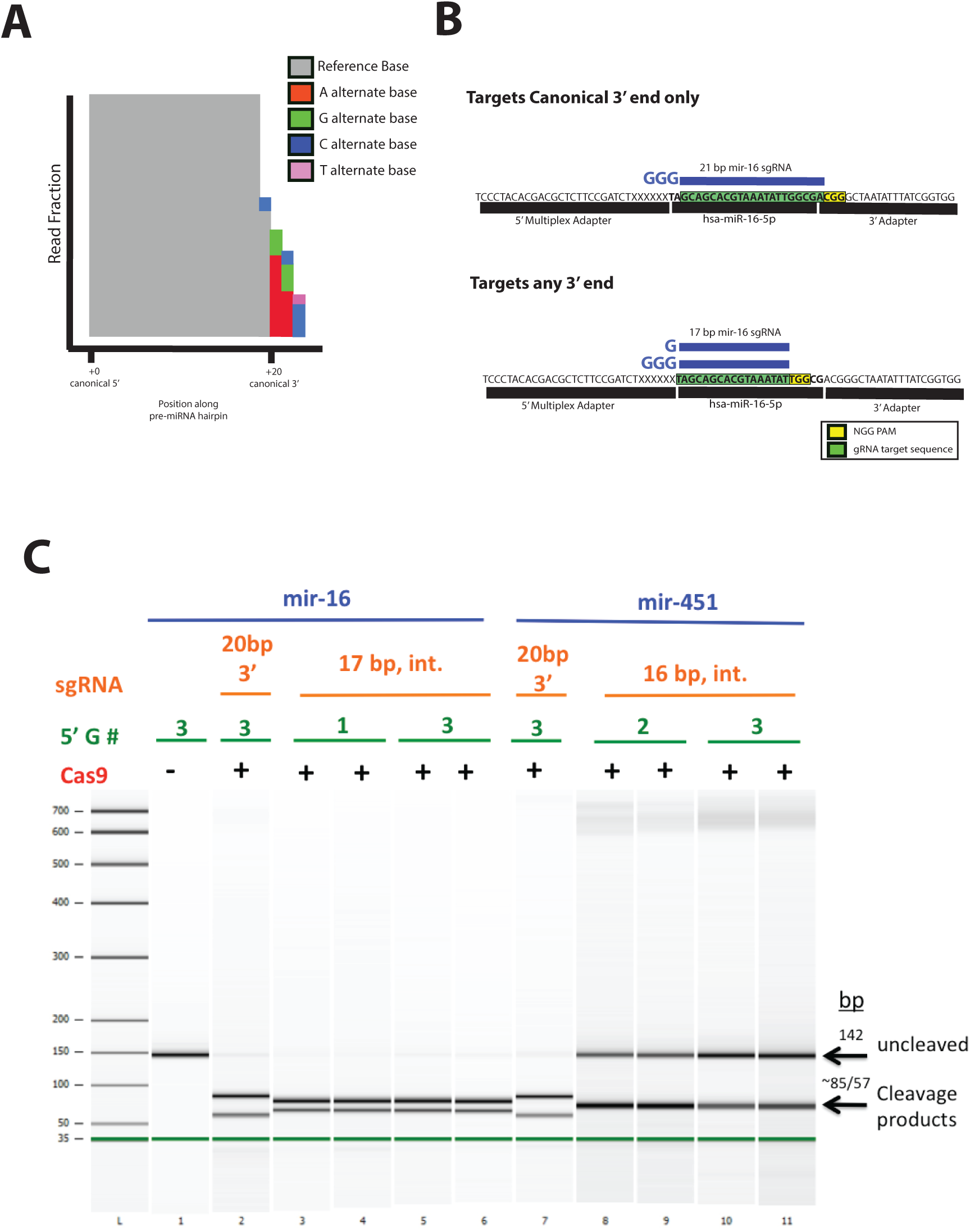
MAD-DASH miRNA targeting with 3’ PAM site or miRNA-internal PAM sites. **A)** Illustration of miRNA 3’ non-templated base addition isoMirs. isomiRs of the same miRNA may not be able to be effectively targeted by an sgRNA designed only to one isomiR with the 3’ Adapter PAM site. **(B)** Sequence level design of MAD-DASH miRNA sgRNAs for 3’ adapter or miRNA-internal PAM use with truncated sgRNAs having varied 5’ Gs. **(C))** Bioanalyzer gel trace for MAD-DASH of miRNA-library gBlocks with design specifications in (B) demonstrates miRNA-PAM efficacy and increasing intolerance of truncated sgRNAs to 5’ mismatches as the sgRNA becomes shorter. Either hsa-miR-16-5p or hsa-miR-451a synthetic library gBlocks were used as substrate for their respective sgRNAs in MAD-DASH reaction. Uncleaved and MAD-DASH cleaved base pair sizes (142 and ~85/57bp (depending on sgRNA), respectively) are indicated. This result supports the work in Fu, et al. 2014 related to the increasing intolerance of truncated sgRNAs to 5’ mismatches as the sgRNA becomes shorter.

**Supplementary Figure S5.**
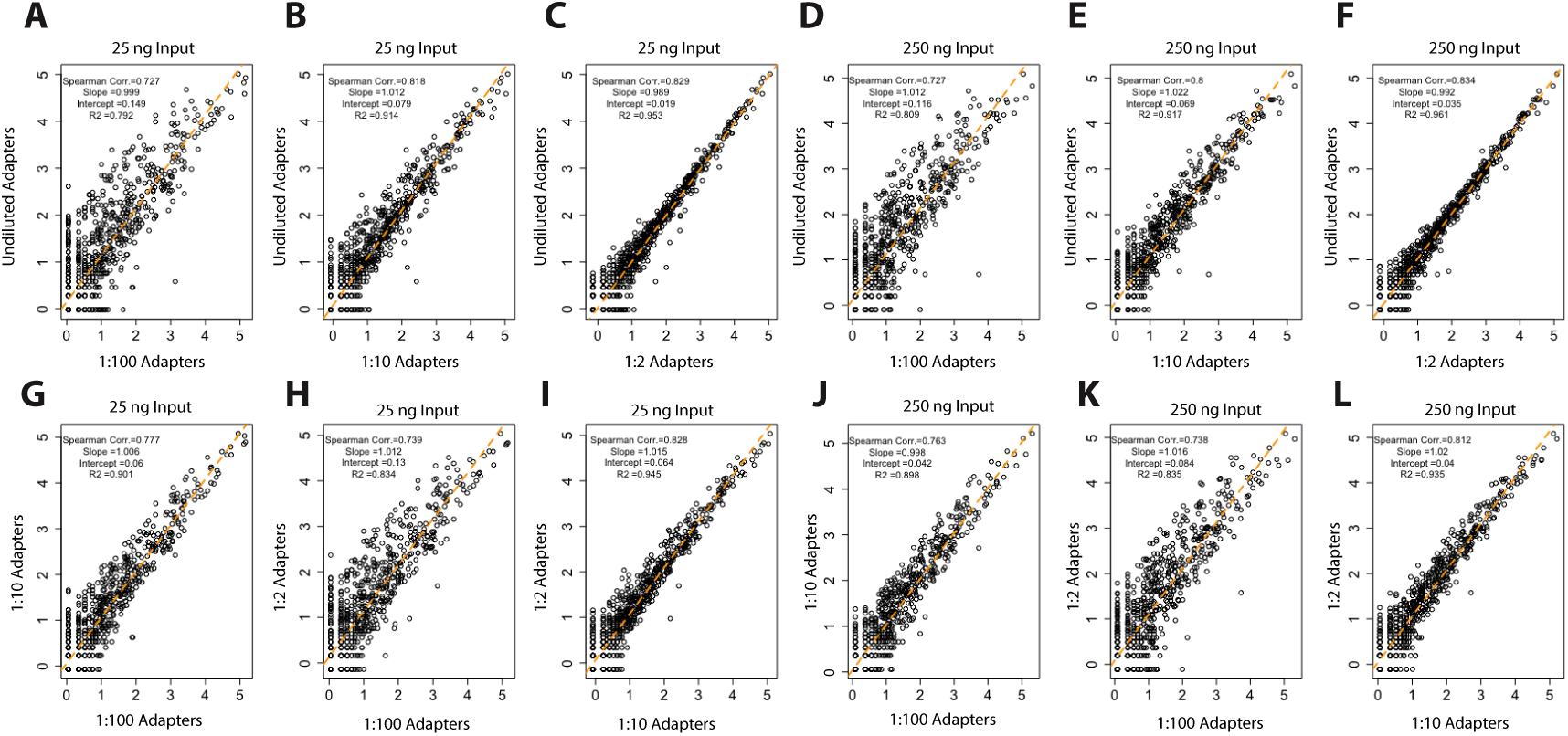
CPM Normalized counts for smRNA-seq libraries constructed from either 25 ng or 250 ng of brain total RNA and varying concentrations of 5’ and 3’ adapters to determine the effect of adapter dilution on library reproducibility. Beginning at the 10 uM / 20 uM 3’- and 5’-adapter concentrations used in our standard protocol, adapters were titrated 1:2, 1:10, and 1:100. As shown for 25 ng (A-C) and 250 ng (D-F) samples, pairwise comparisons between each adapter dilution and our standard concentrations show a concentration-dependent decrease in library correlation. Similarly, pairwise comparisons (G-L) of diluted adapter concentrations to other diluted adapters for both RNA inputs show that libraries made with diluted adapters do not adequately capture RNAs during ligation. This would be expected to significantly impact differential expression measurements between samples and thus we perform our MAD-DASH smRNA-seq protocol with high concentrations of adapters.

**Supplementary Figure S6.**
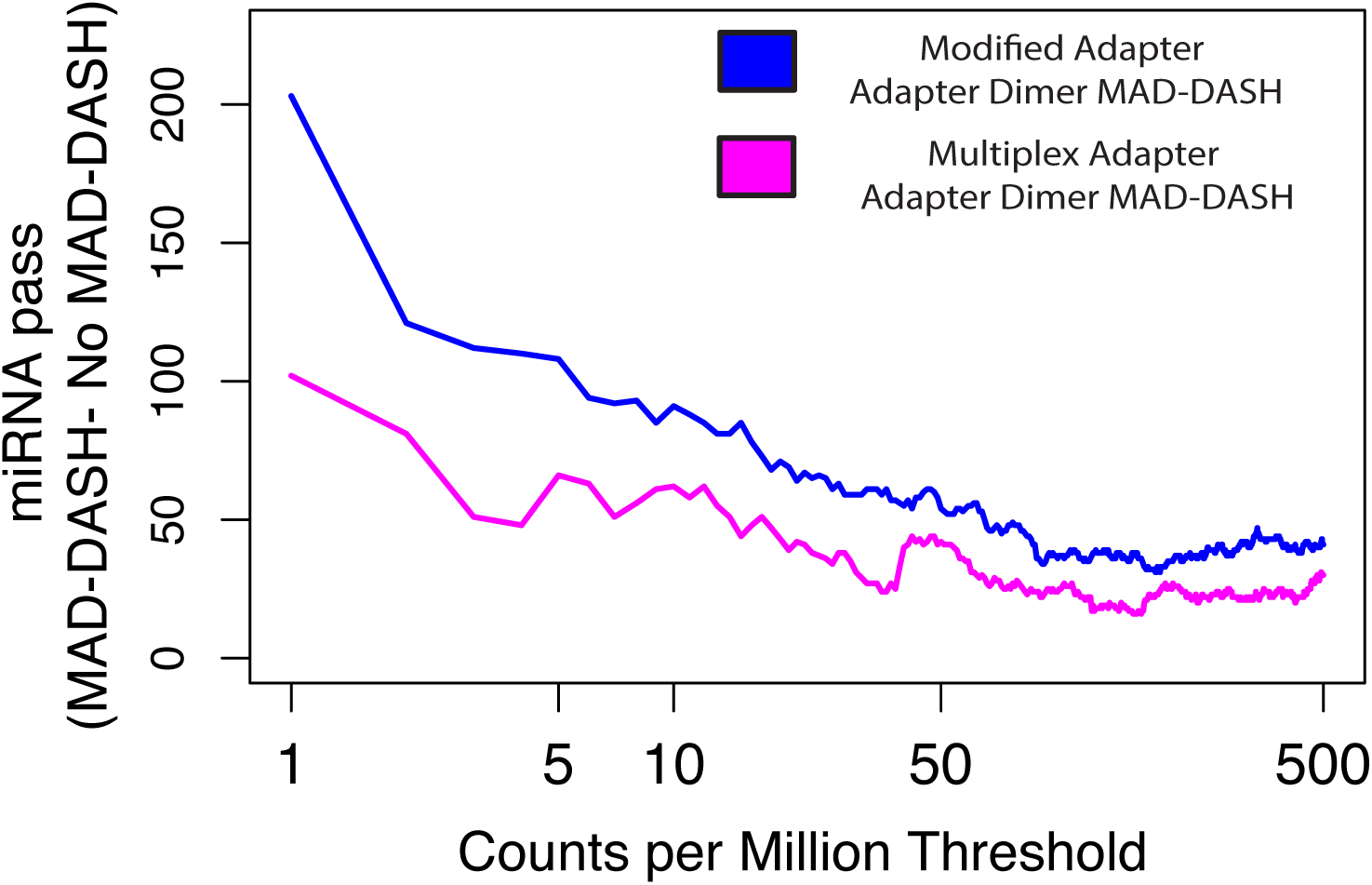
Plot depicting the difference in the number of mature miRNA species reaching a specified count per million threshold between either multiplex or modified adapter dimer MAD-DASH smRNA-seq replicates and their respective no-Cas9/sgRNA MAD-DASH control replicates. Downsampling and CPM normalization was performed as described in Materials and Methods.

**Supplemental Table 1.**
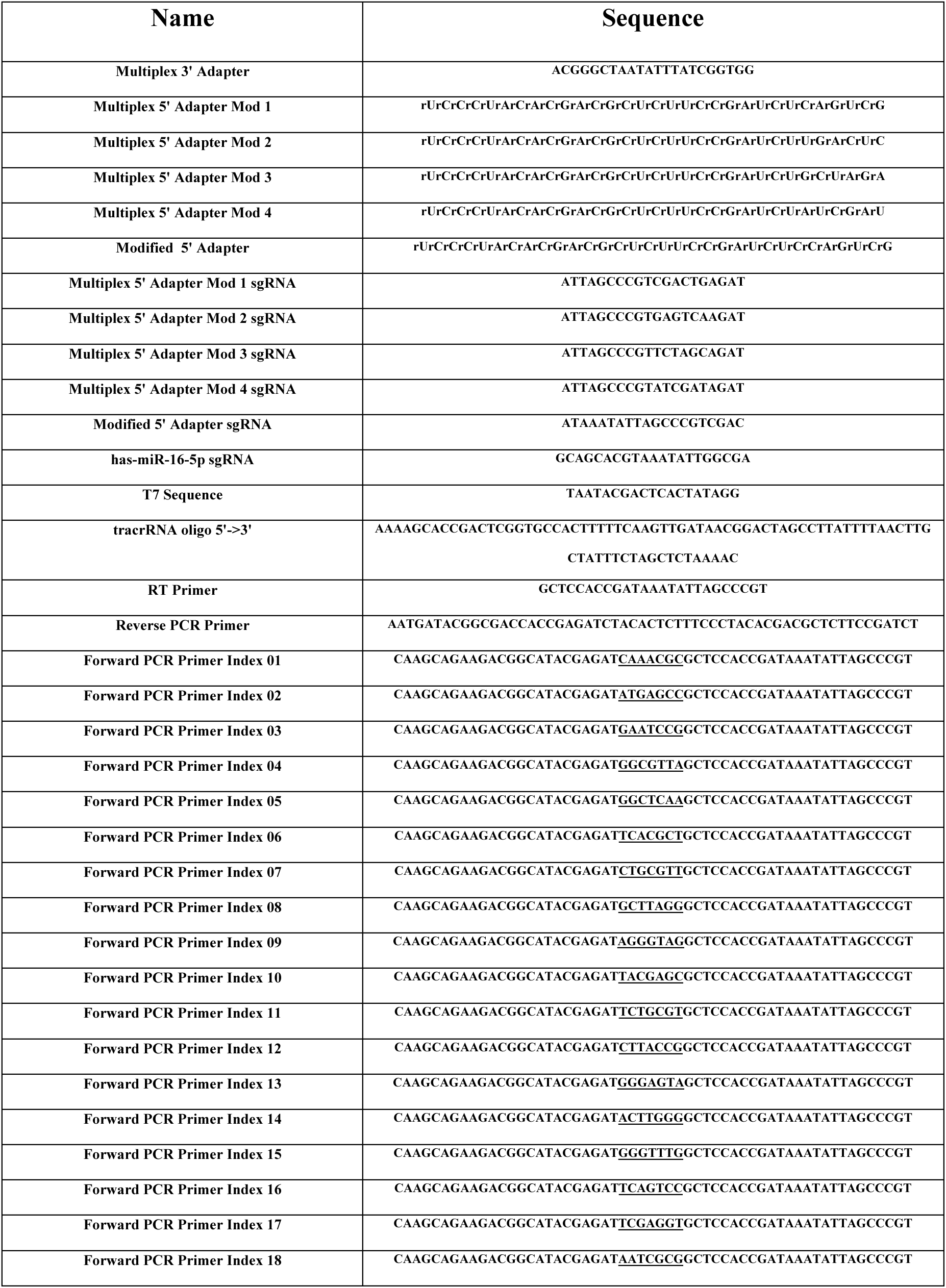
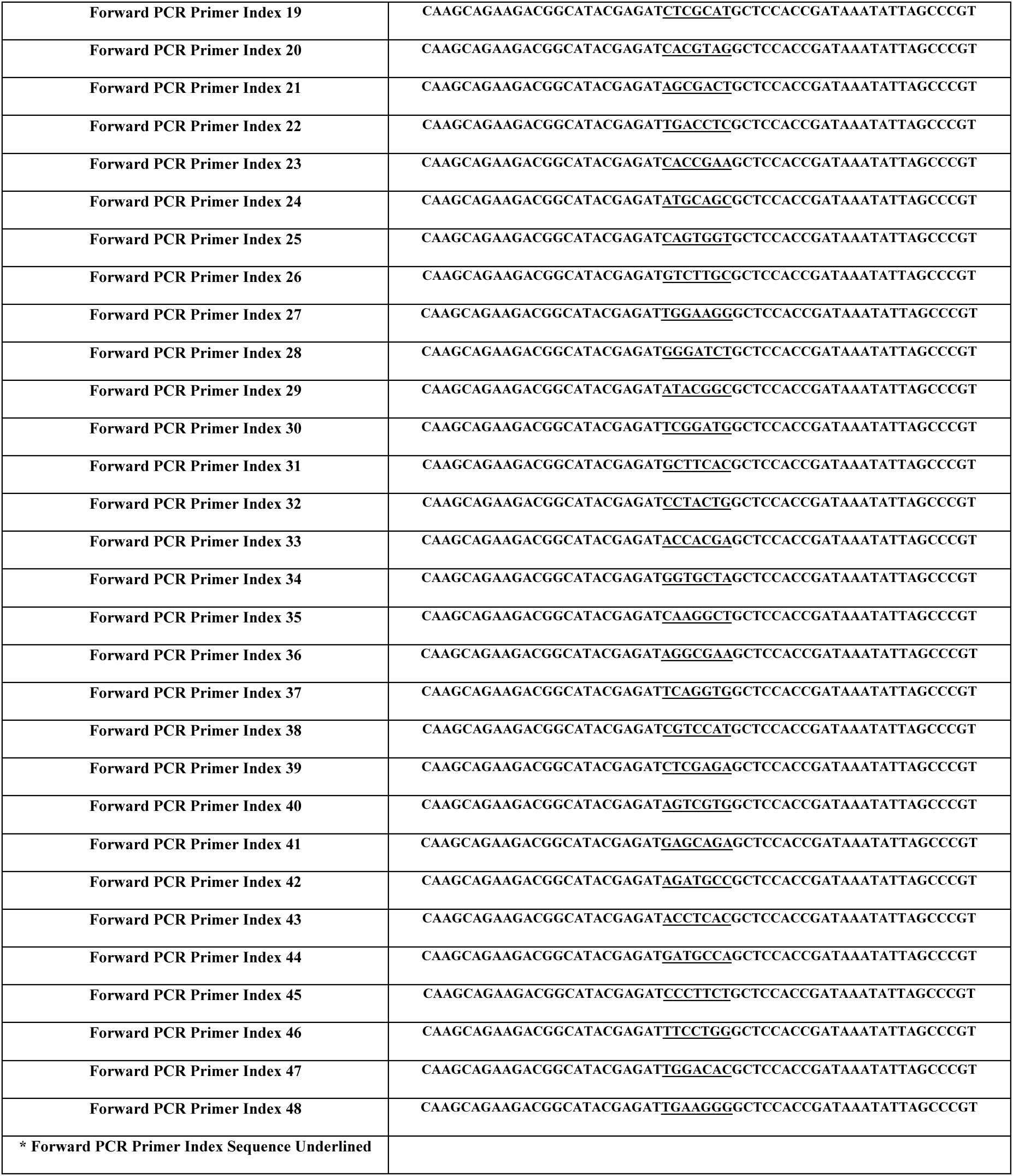
MAD-DASH smRNA-seq Oligo List

**Supplemental Table 2:**
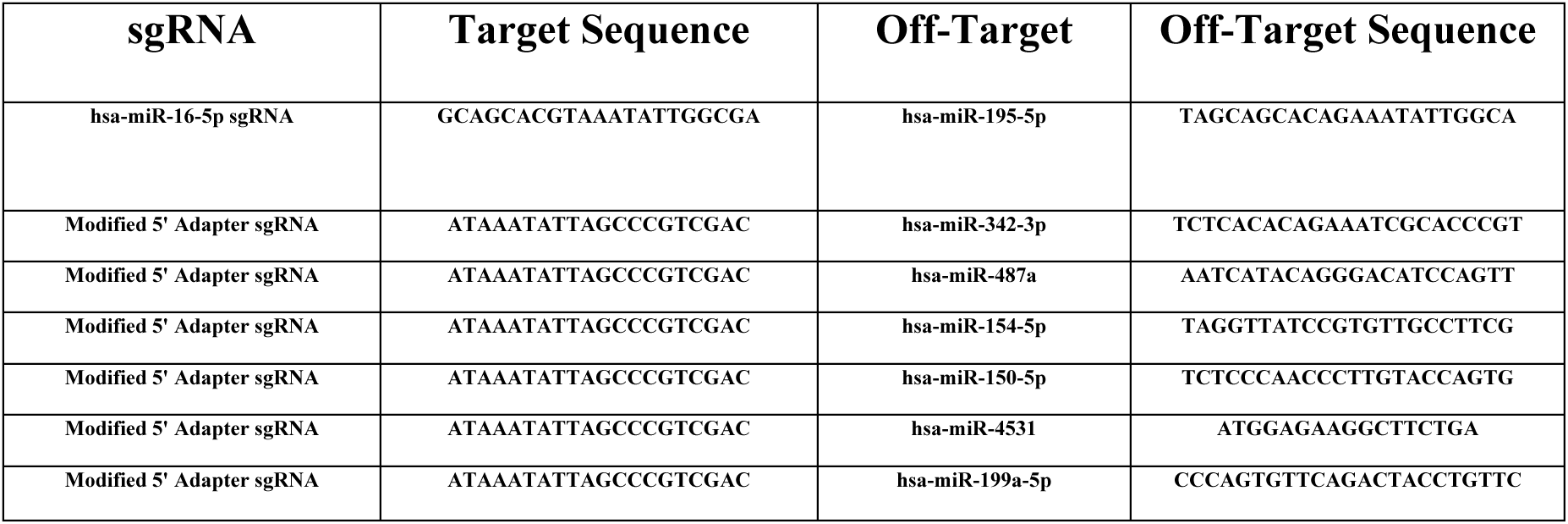
MAD-DASH smRNA-seq significant off-target miRNAs (FDR <0.05, log2FoldChange >1)

**Supplemental Table 3:**
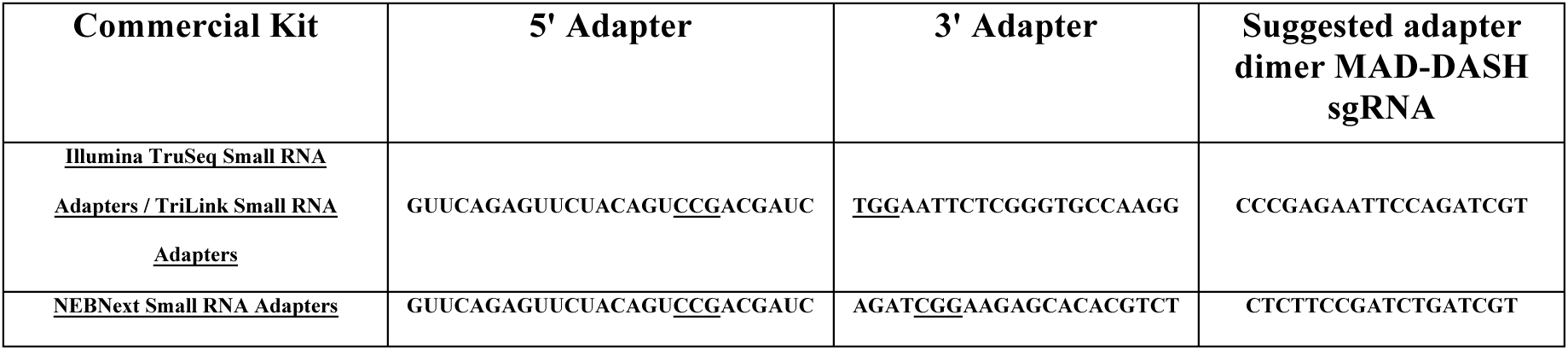
Commercial Small RNA-Seq Adapter Sequences and MAD-DASH Design

